# Immunoglobulins genes in *Neoceratodus forsteri* and *Protopterus annectens* explain the origin of the immunoglobulins of the animals that passed ashore

**DOI:** 10.1101/2022.05.20.492783

**Authors:** Serafin Mirete-Bachiller, Francisco Gambón-Deza

**Affiliations:** Unidad de Inmunología Hospital do Meixoeiro, Vigo, Spain

## Abstract

Sarcopterygii fish have great evolutionary interest since tetrapods and animals that came ashore arose from them. Within immunology, they can teach us about the emergence of Immunoglobulins D, A/X, and Y already present in amphibians. We have studied the genes of the immunoglobulins in the fish Sarcopterygii *Neoceratodus forsteri* and *Protopterus annectens*. In the first fish, we find that several loci for the constant chains of immunoglobulins are distributed in 4 chromosomes. We have found four genes for IgM, a gene for IgW and a gene for IgN. In the second, we find one locus with genes for IgN and IgM and another with one gene for IgW. With these sequences, together with those obtained in other publications, we have been able to study the possible evolution and emergence of immunoglobulin classes. We conclude that there are two evolutionary lines, one focused on IgM and very conservative, and the other focused on IgW, which allows high variability. The W line gave rise to the IgD of 11 domains of reptiles. IgA and IgY are unique since they arose from recombination between the two evolutionary lines. The W line gave origin to the CH1 and CH2 domains, and the M line gave the CH3 and CH4 domains.

## 1. Introduction

The adaptation of vertebrates to land required both favorable external conditions (an increase of oxygen in the terrestrial atmosphere (Hsia et al., 2013)) and previous adaptations to facilitate it (ancestor of bony fishes was already able to breathe air) (Bi et al., 2021). This transition brought with it different genetic innovations that affected locomotion (five digits)(Clack, 2009), anxiolytic ability (Bruce & Neary, 1995) and respiration, initially allowing vertebrates to leave the water temporarily and ultimately permanently (Wang et al., 2021). This transition also had an impact on immunoglobulins (Igs), one of the hallmarks of the adaptive immune system of jawed vertebrates, both in gene organisation and in the appearance of new classes of Igs.

Immunoglobulins are glycoproteins in the gamma globulin fraction with two heavy chains (IgH) and two light chains (IgL). They are linked through disulfide bridges forming a Y-shaped quaternary structure. Immunoglobulin genes appeared 500 million years ago in jawed vertebrates (Flajnik & Kasahara, 2010). Among the current species, the cartilaginous fish are the oldest species. They have these genes(Marchalonis et al., 1998; Clem & Small Jr, 1967). In cartilaginous fish (sharks, skates, and rays), there are three types of Ig: Immunoglobulin M (IgM), immunoglobulin W (IgW) and immunoglobulin NAR (IgNAR) (Kobayashi et al., 1992; Greenberg et al., 1995, 1996). IgM is present in all jawed vertebrates except in *Latimeria chalumnae* (African coelacanth) (Amemiya et al., 2013). IgW is the orthologue of IgD. IgD/W is as old as IgM; therefore, both share a common ancestor that is related to the origin of the immunoglobulins (Ohta & Flajnik, 2006). IgNAR is only present in cartilaginous fish, is a dimer with each chain composed of one V and five C domains, lacks light chains, is homologous to IgW (Roux et al., 1998; Criscitiello et al., 2006). In the animals that came ashore, in addition to IgM and IgD/W, other immunoglobulins emerged, such as IgA/X1, IgA/X2, IgD2 and IgY (Olivieri et al., 2020).

The genes for the immunoglobulins heavy chains (IgH) and light chains (IgL) in Chondrichthyes are in a minilocus organisation. Each minilocus usually consists of a single V, D (IgH only) J, and constant gene segments (Hinds & Litman, 1986). The number of minilocus present varies depending on the type of immunoglobulin and the species (Matz et al., 2020). In contrast, in the rest of the jawed vertebrates, the immunoglobulin genes sequences have a different organisation, so-called translocon (Flajnik & Kasahara, 2010), in which multiple variables (V), diversity (D), and joining (J) segments are upstream of constant (C) domains (Tonegawa, 1983).

A key moment in the evolution of Igs genes is the landfall of vertebrates. Knowledge of the genetic changes that occurred in this process can be given to us by Sarcopterygii fishes (lobe-finned fishes). The genome of *Latimeria chalumnae* (coelacanth), *Neoceratodus forsteri* (giant lungfish) and *Protopterus annectens* (african lungfish) are currently available (Amemiya et al., 2013; Meyer et al., 2021; Wang et al., 2021). The genome of these fish contains a wide variety of transposable-element superfamilies (TEs). The authors point out that the expansion of TEs causes the large size of the genome. TEs contributed to the genomic rearrangement and the appearance of new genes with additional functions or with different regulatory capacities. Tc1 elements (a class of TEs) played a role in the evolution of the vertebrate IgH locus (Magor et al., 1999). Chromosome fusions and fissions contributed to the appearance of genes related to the immune system in *Chiloscyllium plagiosum* (white-spotted bamboo shark) (Zhang et al., 2020).

The authors concluded from the coelacanth genome that the closest living fish to the tetrapod ancestor are lungfishes. Coelacanth immunoglobulin genes have already been studied, finding two genetic loci with V regions and the constant region of the W chain in both. They did not find IgM genes or other Igls (Amemiya et al., 2013). The interest in immunoglobulin lungfish is early. In 1969 two *Neoceratodus forsteri* proteins that corresponded with immunoglobulins were isolated and characterised, one with a higher molecular weight that corresponded with IgM and a second with a lower molecular weight pointed to be typical of these fish (Marchalonis, 1969). Subsequently, a third immunoglobulin of intermediate molecular weight was identified and characterized in *Protopterus aethiopicus* (Litman et al., 1971; Litman, 1975). Other teams have studied messenger RNA sequences of the Protopterus species. In 2003, the nature of this second immunoglobulin was revealed, described as short and long forms of IgW that until then were believed to be restricted to cartilaginous fish (Ota et al., 2003). Later, the study of transcriptomes in two different species of lungfish revealed the presence of two new antibodies, IgN, which must have an evolutionary origin in duplication of IgW and another sequence that they could not completely characterise, that they named IgQ (Zhang et al., 2014).

We have studied the immunoglobulin genes in genomes of *Neoceratodus forsteri* and *Protopterus annectens*. We have found the antibody genes mentioned above and described the gene organisation. Animals that came ashore have a single locus of immunoglobulins. Our results allow an idea of the processes before the single locus structure appeared. The set of sequences allows us to carry out a study that suggests the origin of IgD, IgY and IgA/X genes in amphibians.

## 2. Material and methods

The *Neoceratodus forsteri* genome is in NCBI (assembly neoFor v3). The genome of *Protopterus annectens* is in Chinese Genbank. Due to the large size of the genome, it was studied chromosome by chromosome. It was necessary to split the chromosome to make a blast in some cases. We had a blast locating the genes for immunoglobulins. As a query, we used the published immunoglobulin sequences of *Latimeria chalumnae* (Amemiya et al., 2013) and immunoglobulin sequences from species of protopterus (Zhang et al., 2014). For search exons coding for immunoglobulin CHs in hit segments, the geneid program (Blanco et al., 2007) and tblastx were executed with the queries mentioned above.

Python’s applications were used. The Biopython library to obtain nucleotides and deduced amino acid sequences (Cock et al., 2009). The dna features viewer library was used to create figures (Zulkower & Rosser, 2020). To find the V regions, we use the latest Vgenextractor application (Olivieri et al., 2013).

The alignments of the sequences were made with the Mafft algorithm (Katoh & Standley, 2013). The trees were made with Fasttree (Price et al., 2010) and IQtree (Minh et al., 2020). The Figtree program was used as a viewer (Rambaut, 2007).

## 3. Results

First, we decided to search for the coding exons for the CH regions of the immunoglobulin constant chains and the VH regions. These data give a general information on the number of loscus present in these fish and the classes of immunoglobulins they possess.

### 3.1. Immunoglobulin genes in Neoceratodus forsteri

We looked for immunoglobulin genes using the blast program. Query sequences were already known immunoglobulins from Sarcopterygii fish. We found suggestive sequences in regions of chromosome 7, chromosome 8, chromosome 11 and chromosome 14.

In chromosome 7, we identified one gene for IgW with 11 exons for CHs. In chromosome 8, one gene for IgN with 18 exons for CHs (11 are different and the rest are duplications) and another pseudogene IgN with four exons for CHs (figure 1). In chromosome 9, there are three genes for IgM, two of them pseudogenes. In chromosome 11, there are two additional genes for IgM, and they are pseudogenes. The results indicate the presence of one probable viable gene for IgM, another for IgW and another for IgN.

**Figure 1:**
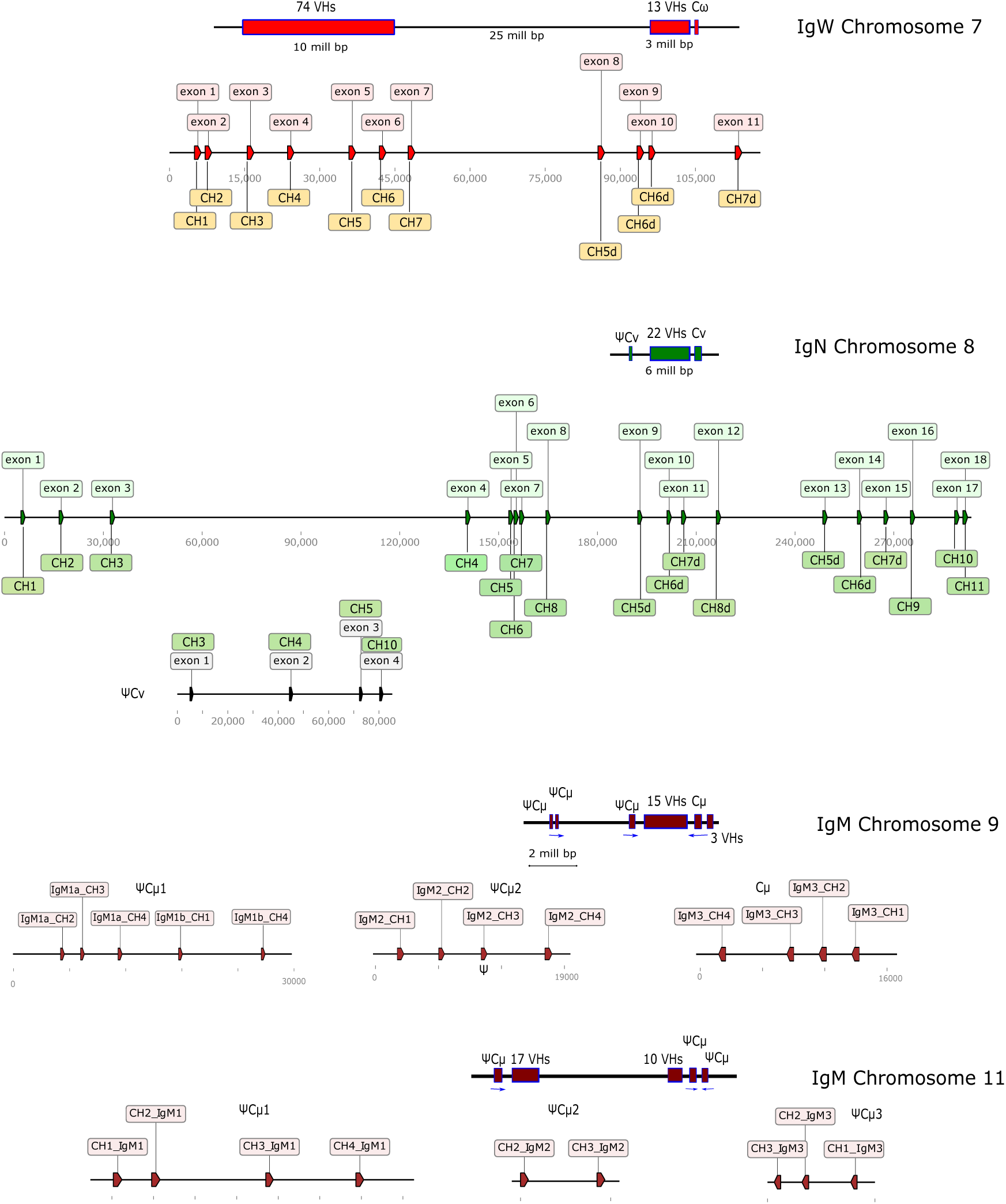
Representation of the coding exons for CHs of immunoglobulins W and N and M in chromosome 7, 8 and 9 respectively. The location of the VH regions in each of the chromosomes is also represented. In chromosome 7 there are 11 exons for IgW, in chromosome 8 there are 28 exons for 2 genes for IgN. 4 belong to one pseudogene and the other 18 exons conform a gene viable. VH regions are presents next to viable gene. In chromosome 9 there are 13 exons for CHS of IgM, 9 in pseudogenes and 4 conform one viable gene. VHs are around of this last gene. The exons have subsequently been labeled to CHs according to the results obtained in the tree in the figure 2.

In the figure 2 we see the sequence relationships between the exons of the genes for immunoglobulins W and N (Also included are the exons found in *P. annectens* that will be explained later). The W gene in *N. forsteri* has seven unique sequences and four duplicates. Those considered unique and duplicates are deduced from the tree in the figure. It is noteworthy that the study of the *Latimeria chalumnae* genome obtained 15 exons for CH domains in this gene and also found seven sequences that can explain the 15. The gene for IgN is complex. At least 11 exons with unique sequences and seven exist seven duplications. The first eight exons plus the last three (exon16, exon17 and exon18) are considered unique.

**Figure 2:**
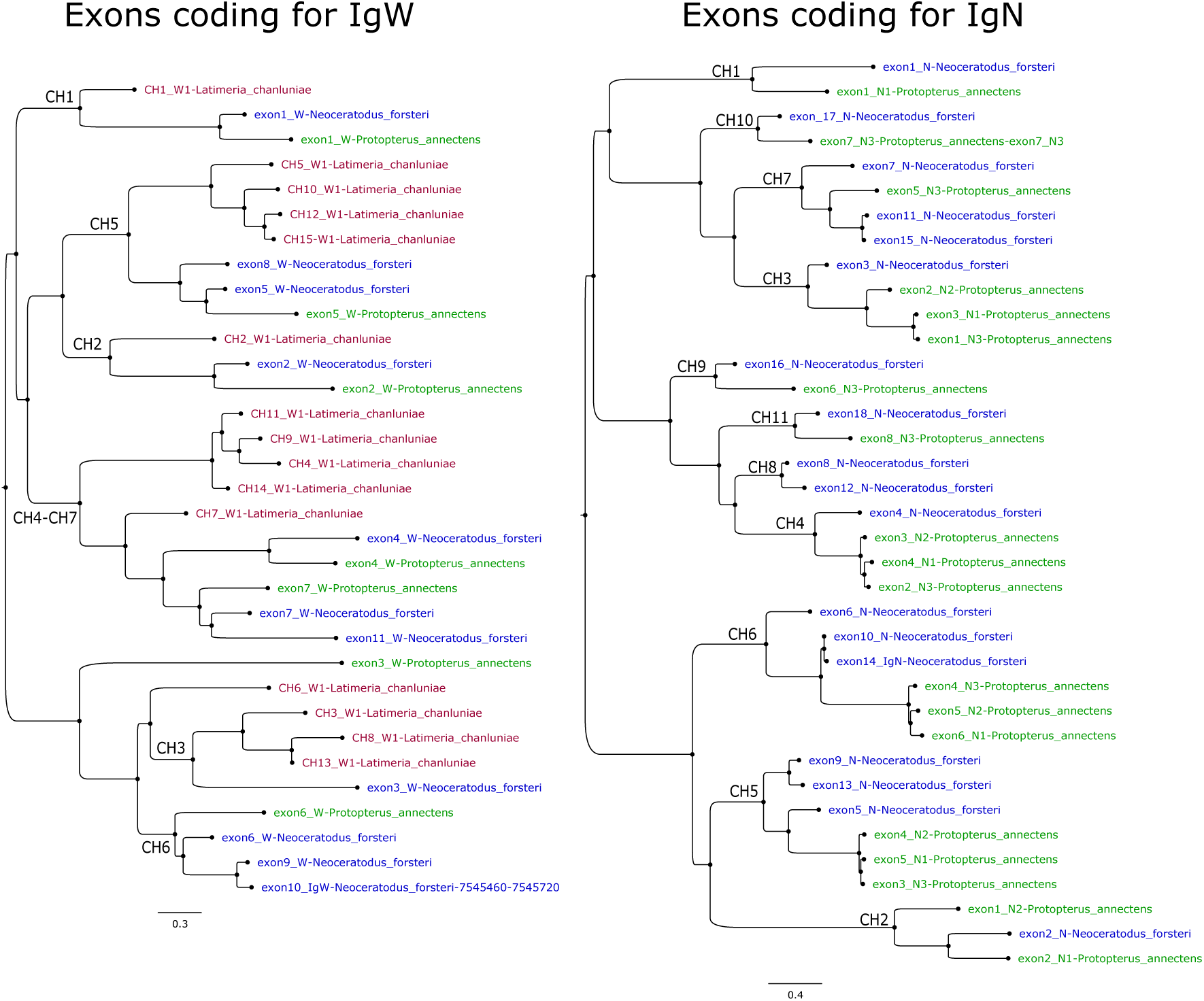
Phylogenetic tree made with the deduced sequences of the coding exons for CHs of the Sarcopterygii species *L. chalumnae, N. forsteri* and *P. annectens*. Each species is in a different color. The purpose of the figure is to deduce the presence of unique CHs and those duplicates for these species. The CH deducted from each clade is noted

In cartilaginous fish, immunoglobulin genes are over several chromosomes, and there is usually a V region for each constant region locus. We search with Vgenenextractor VH regions in *N. forsteri* genome. Most of the VHs regions are on chromosome 7 upstream of the exons for the constant region for IgW (Figure 1). In this chromosome, they form two islands of location. One at the beginning of the chromosome with 74 VH regions distributed in 10 million pairs of basis and another island 25 million downstream from the previous with 13 VH regions. This last island is close to the gene C*ω*. In chromosome 8, there are 22 VH sequences upstream and next to constant region genes for IgN. In chromosome 9, 18 VH regions are grouped next to a gene for IgM. Also, we found 27 VHs regions on chromosome 11 grouped in a region between pseudogenes for IgM

To study the evolutionary relationships in the 4 VH gene locations, we performed an alignment of the deduced amino acid sequences and the phylogenetic tree (figure 3. It is evidenced that the VH of IgW and VH of IgN are in the same clade but different from VHs from IgMs. The results are not absolute, showing some exchanged sequences.

**Figure 3:**
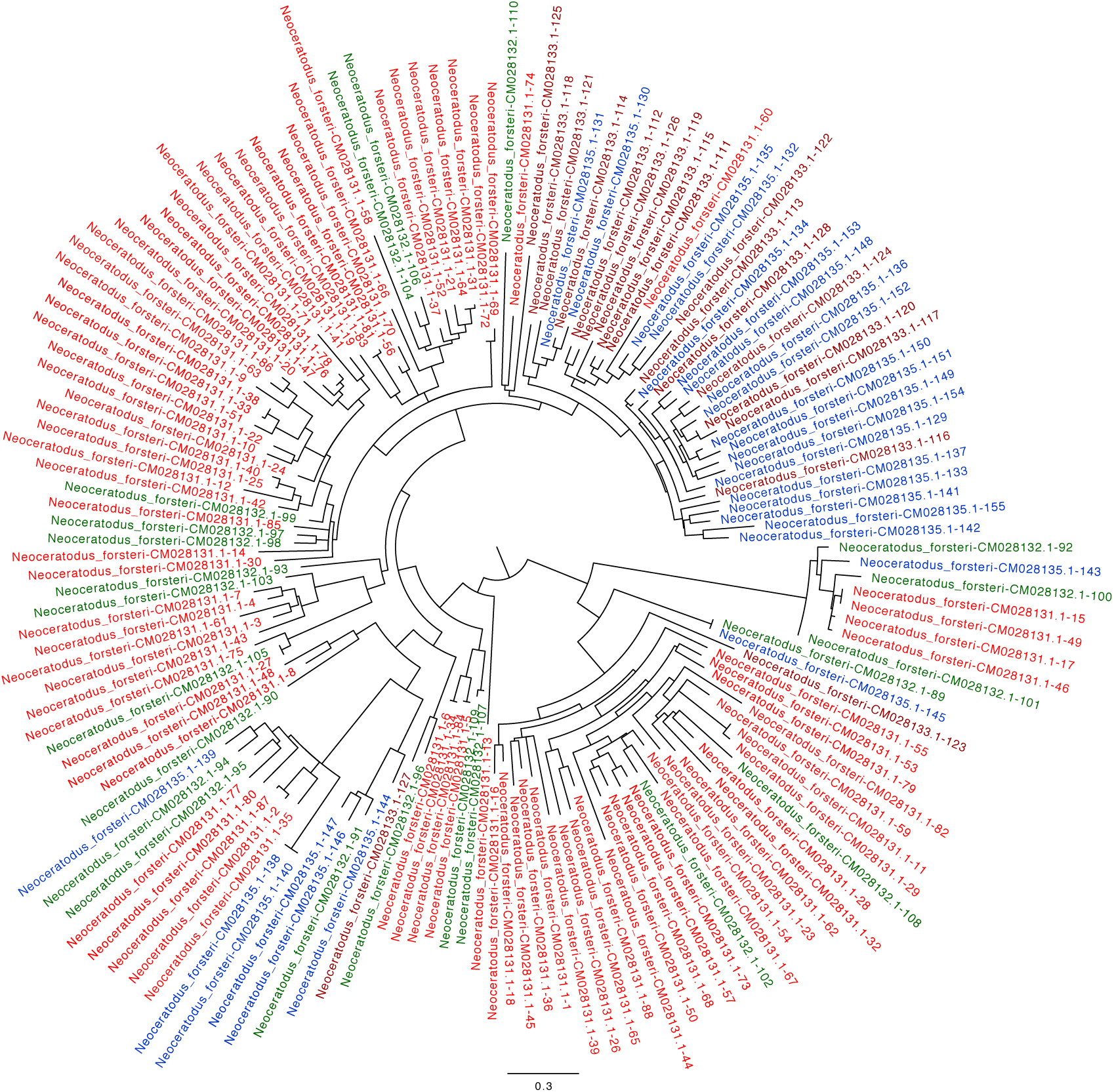
Tree obtained with the amino acid sequences deduced from the sequences of the VH regions found in 4 chromosomes of *N. forsteri*. The colors indicate the location on chromosomes. Red chromosome 7, green chromosome 8, brown chromosome 9 and blue chromosome 11.

### 3.2. Immunoglobulin genes in Protopterus annectens

The blast searches indicated that the genes for the constant chains of immunoglobulins are in chromosome 11 and chromosome 3 (figure 4). In chromosome 11, we find one complex gene for IgN, another for IgW and two genes for IgM. The gene for IgW is far from the cluster of the C*ν* and C*μ* genes. The presence of VH regions is in two zones in chromosome 11. 34 VHs are upstream from the clusters C*ν* and C*μ* and 13 VH upstream next to the C*ω*. In chromosome 3, we find two genes for C*μ*. These genes are relatively close together. The first gene has three VH upstream genes, and the second gene has a single VH.

**Figure 4:**
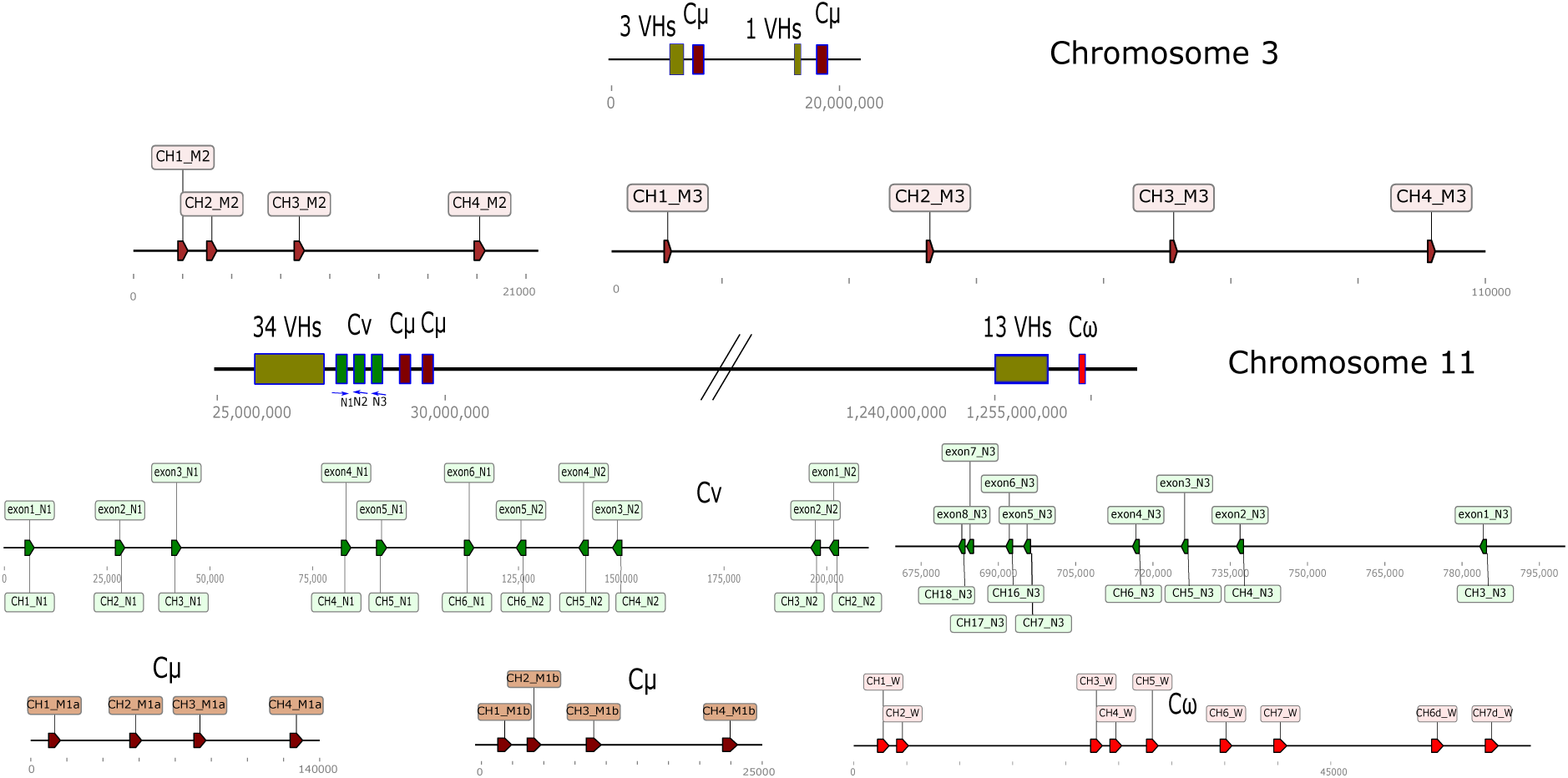
Schematic representation of the immunoglobulin genes found on chromosome 11 of *P. annectens*. The zones with exons C*ν* are in green, C*μ* in brown and C*ω* in red. The C*ν* gene has been divided into three parts as there are exons in chain forward and reverse. The exons have subsequently been labeled to CHs according to the results obtained in the tree in the figure 2.

In a previous study, three different IgM were in the transcriptome of this species (IgM1 KC849709, IgM2 KC849710 and IgM3 KC849711 -REF-). The deduced sequences of the C*μ* genes in chromosome 11 are very similar and correspond to the IgM1 sequences found in the transcriptomes. IgM2 and IgM3 correspond to genes found on chromosome 3.

We also find the sequences described for the hypothetical IgQ (proposed by (Zhang et al., 2014)) in chromosome 16. We find that this sequence comes from two exons. These two exons are in tandem ten times. There are no VH regions in the vicinity or any other data (JH or DH) suggesting an immunoglobulin locus. In conclusion, these sequences do not correspond to an immunoglobulin gene.

The C*ν* gene has 19 exons for CH domains. The gene has exons in the forward chain and in reverse, which determines that we have divided it into three regions. The first region comprises six exons in the forward chain, and they correspond with domains from CH1 to CH6. The second region is in the complementary chain, and there are five exons. These correspond to CH2 to CH6. The third region is also in the reverse chain and has eight exons corresponding to CHs from 3 to 7 plus CH9, CH10 and CH11. The two genes for C*μ* show the classical structure with four domains for CHs. The C*ω* gene has nine exons for CH domains. Duplicate two last exons (C*ω*6 and C*ω*7) are detected in this species as in others previously studied.

The sequence of two IgWs in the transcriptomes is described en protopterus. In *P. Dolloi* both IgW1 and IgW2 are composed of 7 domains. The results of the genomic study indicate that these two immunoglobulins are from the same gene. IgW1 corresponds to the first seven domains, and IgW2 takes the first five and the last two. The CH8 domain has a large region at the beginning rich in cysteines and prolines, suggesting a hinge function.

As mentioned above, there are two regions within chromosome 11 with Vh genes. One region is next to the genes C*ν* and C*μ*s and another next to C*ω*. The sequence relationships between both regions are in the figure 5. Most VH genes around C*ω* are in a separate clade. Regardless of this fact, no evolutionary distance would indicate a divergent evolutionary process.

**Figure 5:**
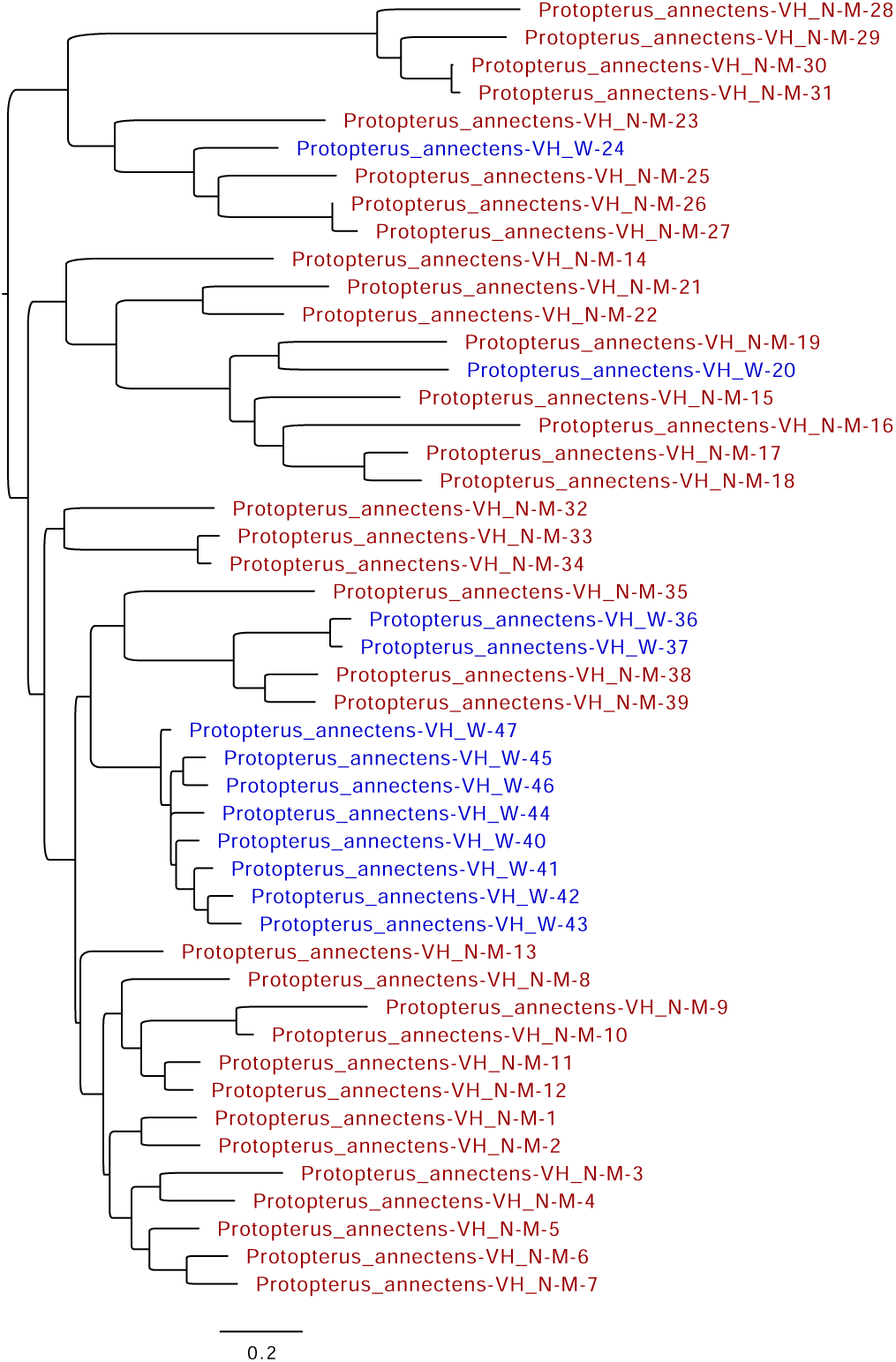
Tree obtained with the amino acid sequences deduced from the sequences of the VH regions found in chromosomes 11 of *P. annectens*. The colors indicate the location on chromosomes. Brown VHs next to C*ν*/C*μ* and blue next to C*ω*.

### 3.3. Evolutive relations of IgN and IgW

We study the evolutive relationship between IgW and the so-called IgN. For this, we obtain the domains to see the relationship between them. These types of antibodies have a large number of domains. Previous publications in Latimeria already described that IgW had seven unique CH domains and later duplications. The IgW described in this article also has seven main domains. To carry out the phylogenetic tree (figure 6), we have taken the domains of IgW and IgN in

**Figure 6:**
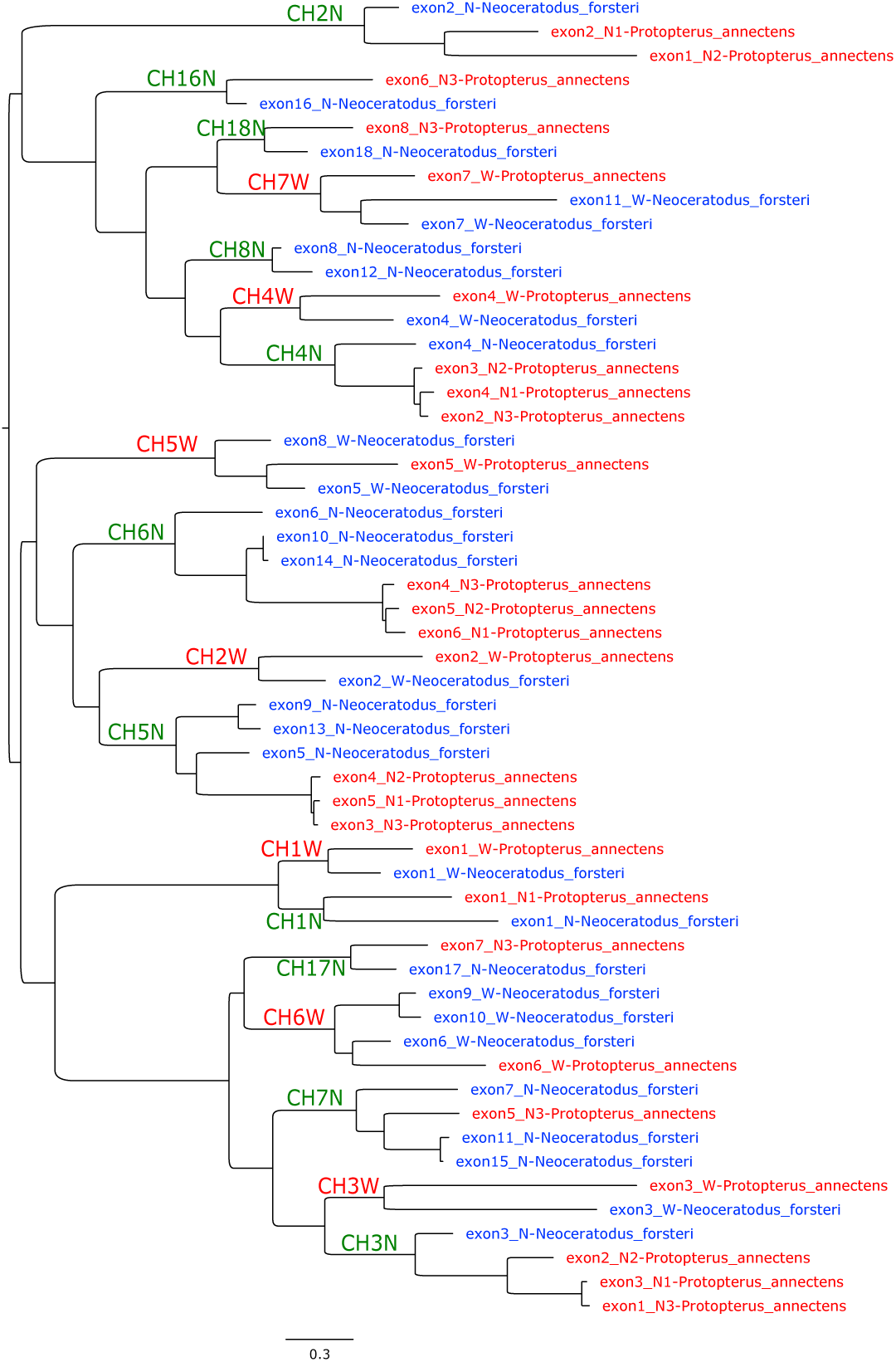
The tree obtained with the amino acid sequences deduced from the sequences of the domains of IgW and IgN from *N. forsteri* and *P. annectens*. Text in blue one specie and red the another. Domains names are in the primary nodes. Nodes of domains of IgW are in red and nodes of IgN in green.

*N. forsteri* and the same domains described in *P. annectens*. The tree shows a correspondence between domains. There are clades with a common origin of most domains indicating that IgN comes from an old duplication of the gene for IgW. However, some exons appear in single clades (CH2N, CH6N, CH16N, CH2W), indicating probable accelerated divergent evolution. (figure 6).

### 3.4. Relationship of Igls of Sarcopterygii with Igls of tetrapods

An old problem in immunology is the origin of Immunoglobulins A and Y. These immunoglobulins are in amphibians (Estevez et al., 2016) but not in species of earlier evolutionary origin. The study of immunoglobulins in sarcopterygian should give knowledge of their origin. To decipher the evolutionary relationships between the immunoglobulins of amphibians, reptiles and sarcopterygians, we made a tree with sequences of CH domains of all the immunoglobulin domains obtained from these species.

The tree has 1265 sequences of domains (Sequences and macro tree in additional information). A scheme of the results of the tree is presented in the figure 7. The unrooted tree image shows four different clades. We take domains IgM as a guide because they are the most conserved immunoglobulin in evolution. In the unrooted tree, each clade is defined by the CH domains of the IgM present. We name each clade as I, II, III and IV (presence of the CH1M, CH2M, CH3M and CH4M domains, respectively). This first observation suggests that all classes of immunoglobulins present today come from one ancestral gene with four domains.

**Figure 7:**
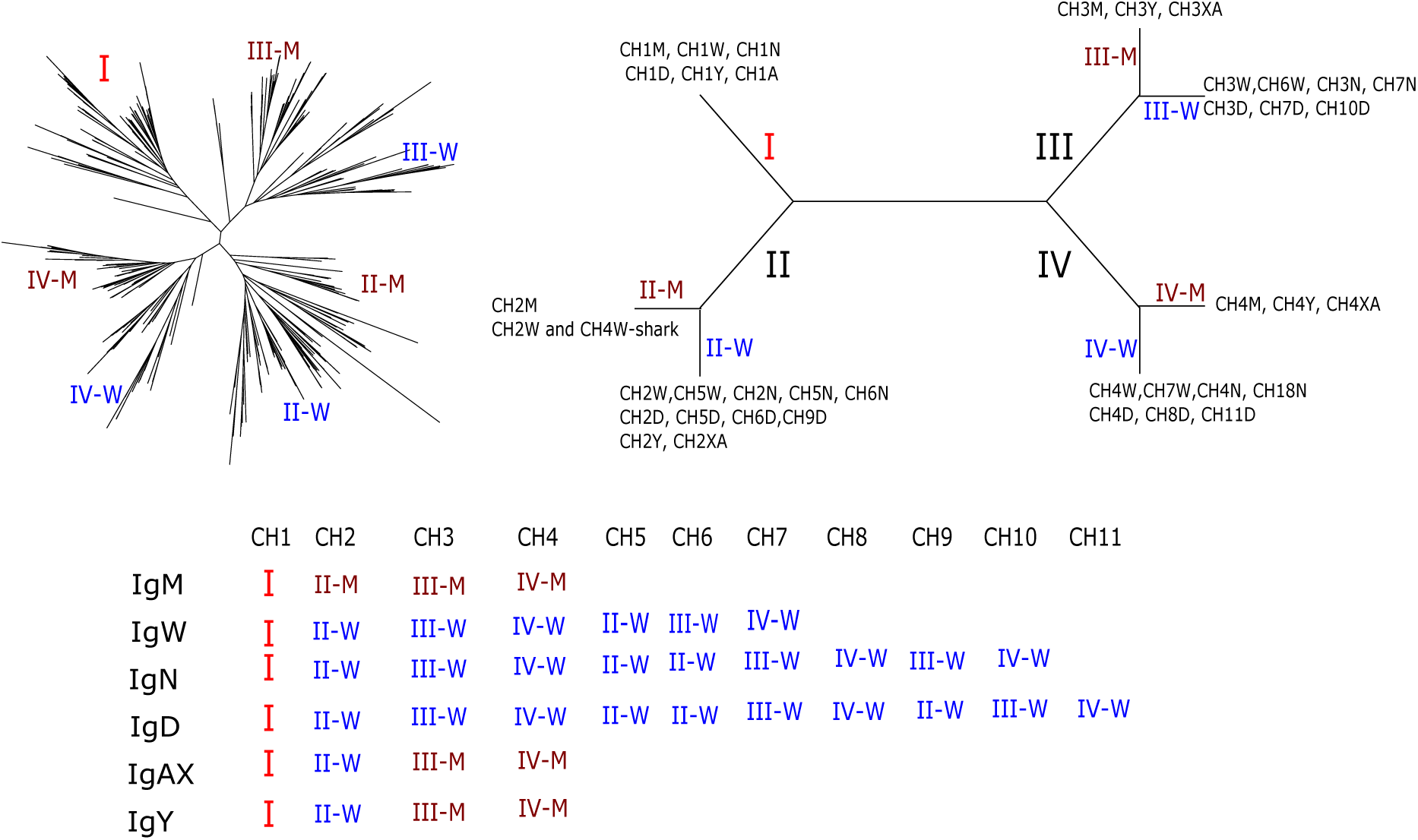
Schematic representation of the results found in a phylogenetic tree with amino acid sequences of the CH domains of immunoglobulins. At the top left is the phylogenetic tree with hundreds of sequences of CHs of immunoglobulins that range from elasmobranchs, sarcopterygian and amphibians. A schematic summary is on the right with the domains found in each clade. In the lower part, the domain composition of each immunoglobulin is expressed.

Clade I exclusively groups all CH1 domains, the sequences of the exons for the CH1 of IgW root the tree. The evolutionary proximity of the clades with sequences for the CH1 of the IgD, IgX/A and IgY stands out.

Clade II has two distinct clades. One of them contains the CH2M domains of all species studied. The other clade presents the rest of the CH2 domains of other immunoglobulins (IgW, IgD, IgY and IgA/X) suggestive of originating from a common ancestor. Also, the sequences for the CH5W, CH5N, CH6N, CH5D and CH10D domains share a common origin). It is interesting to understand the origin of IgY and IgA/X that the CH2 domains of these immunoglobulins are in this subclade.

In Clade III, two different subclades exist. In one are the CH3 domains for IgM, IgY and IgXAs. In the other subclade are IgW, IgN and IgD CH3 domains. These correspond to CH3W, CH6W, CH7N, CH3D, CH7D, and CH10D.

Clade IV, again two subclades are identified. One includes all the CH4M, CH4Y and CH4XA sequences. In the other subclade are several IgW, IgN and IgD sequences.

A diagram with the results of the tree is present in the figure 7. The results indicate that initially, there was a 4-domain antibody that emerged IgM and IgW by duplication. From them, all known tetrapod immunoglobulins originated. IgM is a highly conserved antibody. IgW has undergone many evolutionary changes in the number of domains accepting elongation by terminal duplicate domains. Furthermore, the IgW will give the IgD of 11 amphibians and reptilians.

The results also suggest the origin of the immunoglobulins IgY and IgA/X. These immunoglobulins arose from the recombination of the W line with the M line. In clade I, the presence of the W and M lines cannot be discerned since there has been an exchange of CH1 between different immunoglobulins. The CH2 comes from the IgW line, and the CH3 and CH4 domains from the IgM line. The results are suggestive that its origin was due to recombination between the two main lines. In the lower part of the figure 7 the evolutionary origin of the domains that make up the main classes of immunoglobulins is represented. It stands out that IgY and IgA/X present evolutionary domains of both IgW and IgM.

## 4. Discussion

The present results of the distribution of the genes for the immunoglobulin heavy chain in the two species of Sarcopterigii fish studied allow us to understand the processes that occurred before the appearance of immunoglobulins in the tetrapods. Our results show that *N. forsteri* still presents multiple individualized loci for each of the immunoglobulins. Clusters VH exist next to genes for constant regions. In addition, there are pseudogenes for IgM. The *P. annectens* genome is in a more evolved state. There are only two loci in the same chromosome, although very far apart. Furthermore, for the first time, a locus with genes for two different classes of immunoglobulins (IgM and IgN) has been identified, which can be considered the prelude to a unique locus of immunoglobulins later found in amphibians. In both tetrapods and fish, we find the translocon constituted. In sarcopterygian, we find an intermediate situation. They already have clusters of VH genes and a decrease in immunoglobulin loci. The existence of C*μ* pseudogenes supports the hypothesis of evolutionary pressure for creating a single locus.

The organization of immunoglobulin genes in the sarcopterygian indicates an evolutionary trend toward creating the translocon. In elasmobranchs, immunoglobulin genes are on several chromosomes. In Actinopterigian and tetrapods, the immunoglobulin locus is usually unique. It is probable that grouping the genes was pressed by the need to obtain specific responses, so one cell antibody’s pathway was favoured (Matz et al., 2021).

One of the classic problems of immunology is the origin of the classes of immunoglobulins present today in tetrapods. The passage of animals to land led to new immunoglobulins classes and the ability to switch in immunoglobulin classes. Data indicate that the sarcopterygian was the fish that gave rise to the tetrapods. Genome sequences of two species and RNA sequences of others allow the search for ancestors of immunoglobulins from terrestrial animals. We obtained sequences that allow us to make an approximation to understand the origin of antibodies in terrestrial animals. In amphibians, several IgDs have been described, which are related to IgDs of reptiles and mammals. The 11 domains that explain all IgDs from terrestrial animals (Olivieri et al., 2021; Estevez et al., 2016) can now be explained by deducing their origin from IgW/IgN. All the domains are related to some of the line W.The IgD in amphibians and reptiles must be derived from an IgN gene. Within the phylogenetic tree, the IgD domains are closer to those of IgN. In addition, as can be seen in the diagram of the figure 7, the structure of IgN is very similar to that of IgD with 11 domains. It also supports the fact that in the genome of *P. annectens* the gene for IgN is found together with the IgM genes.

The origin of IgA was proposed to come from the recombination of IgM and IgY (2 first domains of IgY and the two terminals of IgM (Deza et al., 2007)). In this study, the recombination origin of IgA is confirmed with the CH3 and CH4 domains from IgM. The origin of the first two domains does not come from IgY but the same evolutionary line as the first two domains of IgY. Regardless, the results obtained indicate that these two antibodies come from the recombination of the M line with the W line and differentiate them from the other antibodies. It is noteworthy that the IgD2 found in reptiles (Gambón-Deza & Espinel, 2008)also has a recombination origin, as expressed here, therefore resembling these antibodies.

The changes that occurred up to the conformation of the immunoglobulin genes in tetrapods are expressed in the figure 8. In Sarcopterigii fish, intermediate structures of immunoglobulin genome organization are conserved until now. This organization helps to understand how the multilocus formation present in elasmobranchs evolved to the formation of the traslocon, the generation of a single locus and with several genes for constant region present in tetrapods.

**Figure 8:**
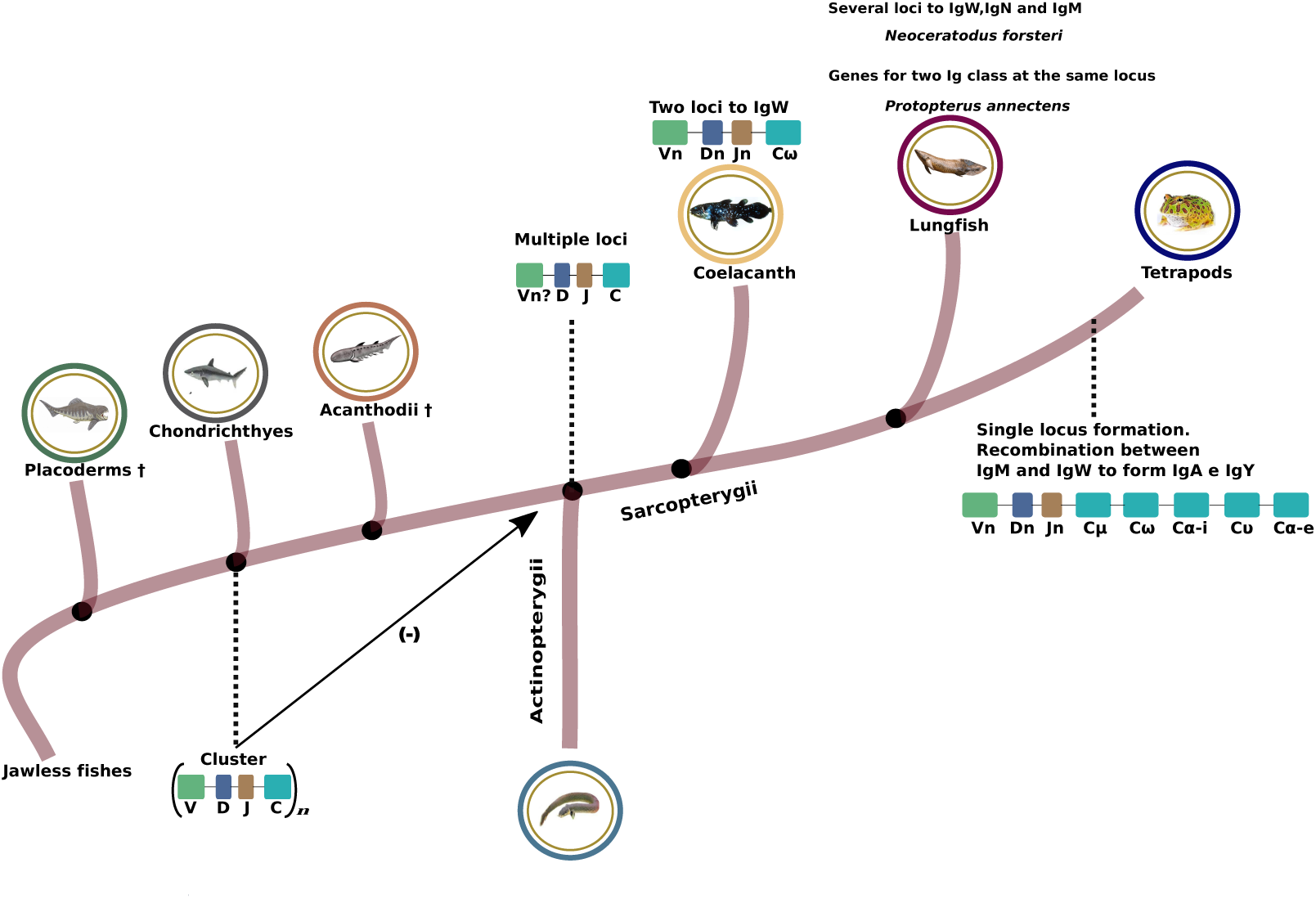
Schematic representation of the changes that occurred in the genome organization of immunoglobulin genes from elasmobranchs to Tetrapods. elasmobranchs present multiple loci with gene units composed of one V gene, D and J mini-genes, and a gene for constant chain (can be for IgM or IgW). In some loci, there was a multiplication of the V genes. In this way, the translocon was formed in which a V had to be selected. This process must have occurred before the divergence between Sarcopterigii and Actinopterygii. In Sarcopterigii fish, the number of loci decreases and some loci already appear with the presence of two genes for the constant regions. Later, in the tetrapods that reached land, only a single locus with multiple V genes and several genes for the constant region was detected.

## Supporting information

suplemental sequences

suplemental tree

## 6. Additional information

